# Modelling the Effects of Ongoing Alpha Activity on Visual Perception: The Oscillation-Based Probability of Response

**DOI:** 10.1101/752766

**Authors:** Agnese Zazio, Marco Schreiber, Carlo Miniussi, Marta Bortoletto

**Author notes:** **Corresponding author:** Agnese Zazio, Cognitive Neuroscience Section, IRCCS Istituto Centro San Giovanni di Dio Fatebenefratelli, Via Pilastroni 4, 25125 Brescia (Italy). **HIGHLIGHTS** - A novel, interdisciplinary model to link prestimulus oscillations and perception - Alpha-gamma cross-frequency predicts probability of response to visual stimuli - Different meso-scale neural mechanisms selectively affect the psychometric function.

## Abstract

Substantial evidence has shown that ongoing neural activity significantly contributes to how the brain responds to upcoming stimuli. In visual perception, a considerable portion of trial-to-trial variability can be accounted for by prestimulus magneto/electroencephalographic (M/EEG) alpha oscillations, which play an inhibitory function by means of cross-frequency interactions with gamma-band oscillations. Despite the fundamental theories on the role of oscillations in perception and cognition, a clear theorization of the neural mechanisms underlying prestimulus activity effects that includes electrophysiological phenomena at different scales (e.g., local field potentials and macro-scale M/EEG) is still missing. Here, we present a model called the oscillation-based probability of response (OPR), which directly assesses the link between meso-scale neural mechanisms, macro-scale M/EEG, and behavioural outcome. The OPR model includes distinct meso-scale mechanisms through which alpha oscillations modulate M/EEG gamma activity, namely, by decreasing *a)* the amplitude and/or *b)* the degree of neural synchronization of gamma oscillations. Crucially, the OPR model makes specific predictions on the effects of these mechanisms on visual perception, as assessed through the psychometric function.

**SIGNIFICANCE STATEMENT:** The oscillation-based probability of response (OPR) is grounded on a psychophysical approach focusing on the psychometric function estimation and may be highly informative in the study of ongoing brain activity because it provides a tool for distinguishing different neural mechanisms of alpha-driven modulation of sensory processing.

## INTRODUCTION

The response of neurons is well known to not merely depend on external input; indeed, a repeated presentation of the same stimulus gives rise to highly variable responses at the neural level as well as at the behavioural level (Arieli et al., 1996; Vogels et al., 1989). Interestingly, such response variability can be accounted for by fluctuations in ongoing brain activity, as revealed by invasive (Arieli et al., 1996) and non-invasive (Kayser et al., 2016; Weisz et al., 2014) electrophysiological recordings, functional magnetic resonance imaging (Baldassarre et al., 2012; Hesselmann et al., 2008) and behavioural (Song et al., 2014) studies.

The time-frequency pattern of brain oscillations represents a key feature by which ongoing activity shapes both neural and behavioural responses (Ai and Ro, 2014; Baumgarten et al., 2016; Haegens et al., 2011; Kayser et al., 2016; Leske et al., 2015; Linkenkaer-Hansen, 2004; Mazaheri et al., 2009; Schubert et al., 2008; van Dijk et al., 2008). Specifically, ongoing oscillations within the alpha band (frequency range: 8-13 Hz), as measured by magneto/electroencephalographic (M/EEG) recordings, represent a major factor in accounting for response variability in the domain of visual perception (Busch et al., 2009; Iemi et al., 2017; Iemi and Busch, 2018; Lange et al., 2014; Mathewson et al., 2009; H. van Dijk et al., 2008).

In recent decades, neuroscientists have developed fundamental theories highlighting the role of oscillations in brain dynamics, such as in neural inhibition (inhibition timing hypothesis, Klimesch et al., 2007; gating by inhibition, Jensen and Mazaheri, 2010) and neural communication (communication through coherence, Fries, 2005). Nevertheless, the micro- (e.g., single-unit measurements) and meso-scale (e.g., local field potential, LFP) neural mechanisms that contribute to M/EEG oscillatory activity and their link with behaviour are far from established (Cohen, 2017; Musall et al., 2014).

## THE OSCILLATION-BASED PROBABILITY OF RESPONSE

The oscillation-based probability of response (OPR) is a model that fuses neurophysiological evidence, mathematical modelling and a psychophysical approach. The aim of the model is twofold: first, to identify potential neural mechanisms of alpha-gamma interactions that support the effects of prestimulus M/EEG oscillations on visual perception (Lange et al., 2014; Ruhnau et al., 2014; Sadaghiani and Kleinschmidt, 2016; Zoefel and VanRullen, 2017); second, to provide a framework with clear predictions on the effects on perception, so that behavioural performance can be exploited to distinguish the underlying neural mechanisms that cannot be disentangled by means of M/EEG.

The OPR is based on the following key points: 1) alpha oscillations affect visual perception; 2) alpha activity plays an inhibitory role; 3) alpha inhibition occurs through alpha-gamma cross-frequency interactions; 4) alpha-modulated gamma oscillations affect the probability of neurons in sensory areas to respond to an incoming stimulus; and 5) the probability of response is selectively modulated by distinct neural mechanisms of alpha inhibition.

Crucially, the OPR includes the estimation of the response probability for a wide range of stimulus intensities and therefore generates a psychometric function. In this framework, the estimated psychometric function represents an extremely powerful tool because it provides suggestions on the selective involvement of different neural mechanisms, which cannot be discerned non-invasively (Cohen, 2017; Whittingstall and Logothetis, 2009). We therefore suggest that the OPR could be applied to make hypotheses on distinct mechanisms involved in alpha-driven modulation of sensory processing, based on selective modification of the psychometric function.

## 1. ALPHA OSCILLATIONS AFFECT VISUAL PERCEPTION

Oscillatory activity within the alpha band is dominant in the human brain; it represents the strongest electrophysiological signal that can be measured non-invasively and is observed transversely across cognitive domains (Berger, 1929; Klimesch, 2012).

The effect of spontaneous variations in M/EEG alpha activity on perception is well documented. Typically, to detect the effects of such spontaneous fluctuations, studies employ stimuli at the individual sensory threshold, i.e., the so-called “near-threshold stimuli”, by definition detected in half of the trials (for a review see Ruhnau et al., 2014). Single-trial analysis is then performed to assess within-subjects variance (Pernet et al., 2011).

Experimental evidence has shown that perceptual performance in the visual domain is affected by prestimulus M/EEG alpha power. Specifically, trials with low alpha power preceding stimulus onset lead to a higher probability of stimulus detection, both between and within subjects (**Figure 1A**; Busch and Van Rullen, 2010; Hanslmayr et al., 2007; van Dijk et al., 2008), and recently, this effect has been suggested to potentially not be due to an improved perceptual acuity but rather to a more liberal criterion in the response (Iemi et al., 2017; Limbach and Corballis, 2016). Furthermore, a few studies reported that the visual detection rate differs between opposite phases of 7-10-Hz oscillations (**Figure 1B**; Busch et al., 2009; Busch and Van Rullen, 2010; Mathewson et al., 2009; but see Benwell et al., 2017; Benwell et al., 2013). Consistently, ongoing M/EEG alpha power and phase have been shown to be relevant factors in predicting transcranial magnetic stimulation-induced phosphene perception (Dugué et al., 2011; Romei et al., 2008).

**Figure 1.**
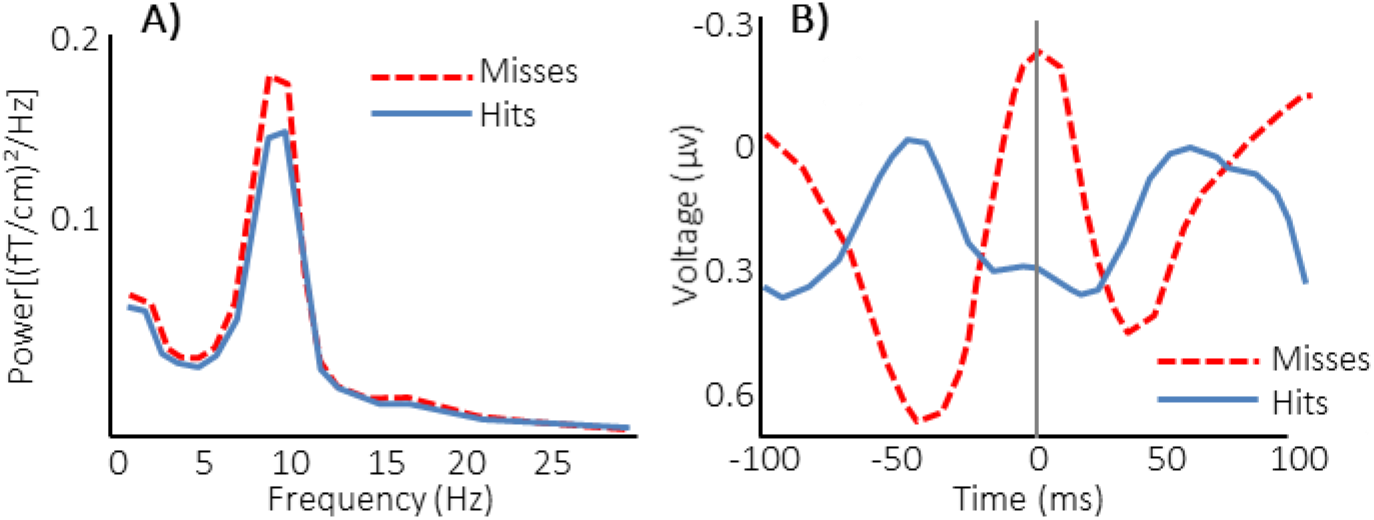
Effects of prestimulus M/EEG alpha oscillations on visual detection. **A)** Lower alpha power preceding the stimulus leads to a higher proportion of hits (grand average of the spectra calculated in the second before target onset in occipital MEG channels; adapted from van Dijk et al., 2008). **B)** Opposite alpha phase at target onset (time: 0 ms) for hits and misses (grand average event-related potential at EEG channel Pz; adapted from Mathewson et al., 2009).

Moreover, a few studies have provided evidence of a causal role of prestimulus alpha oscillations on perception by modulating the alpha rhythm. A traditional way to modulate alpha power is by means of spatial attention: alpha power is lower for the attended than for the unattended visual hemifield (Worden et al., 2000; Sauseng et al., 2005), thereby affecting detection performance (Busch and Van Rullen, 2010). More recently, alpha oscillations have been modulated with entrainment mechanisms, reflecting the phase alignment of the brain’s oscillatory activity to external rhythmic stimulation by sensory stimulation (e.g., visual or auditory rhythmic stimuli; Spaak et al., 2014; Henry and Obleser 2012) and through non-invasive brain stimulation techniques (Thut et al., 2017; Thut et al., 2011; Helfrich et al., 2016). Entrainment of an endogenous alpha rhythm results in enhanced power and phase-locking, and, in turn, the entrained alpha power and phase modulate perception, consistent with findings on the effects of spontaneous oscillations (Landau and Fries, 2012; Romei et al., 2010; Spaak et al., 2014).

In short, there is both correlational and causal evidence that ongoing alpha oscillations shape visual perception.

## 2. ALPHA ACTIVITY PLAYS AN INHIBITORY ROLE

The direction of behavioural effects of alpha activity described in the literature, i.e., the larger the alpha power, the lower the probability of perceiving a visual stimulus, have led to the most widespread interpretation of considering alpha activity as playing an inhibitory role (Klimesch et al., 2007; Jensen and Mazaheri, 2010; but see Palva and Palva, 2007).

According to the gating by inhibition (GBI; Jensen and Mazaheri, 2010) hypothesis, alpha inhibition appears to be cyclic and linked to the phase of alpha oscillations; rhythmic alpha activity serves as a pulsed inhibition that modulates the time window for sensory processing, i.e., the duty cycle. In this view, when alpha power is high (i.e., high inhibition), the duty cycle is shorter than that when alpha power is low (Jensen & Mazaheri, 2010; Mathewson et al., 2009), as shown by the green trace in **Figure 2A**.

**Figure 2.**
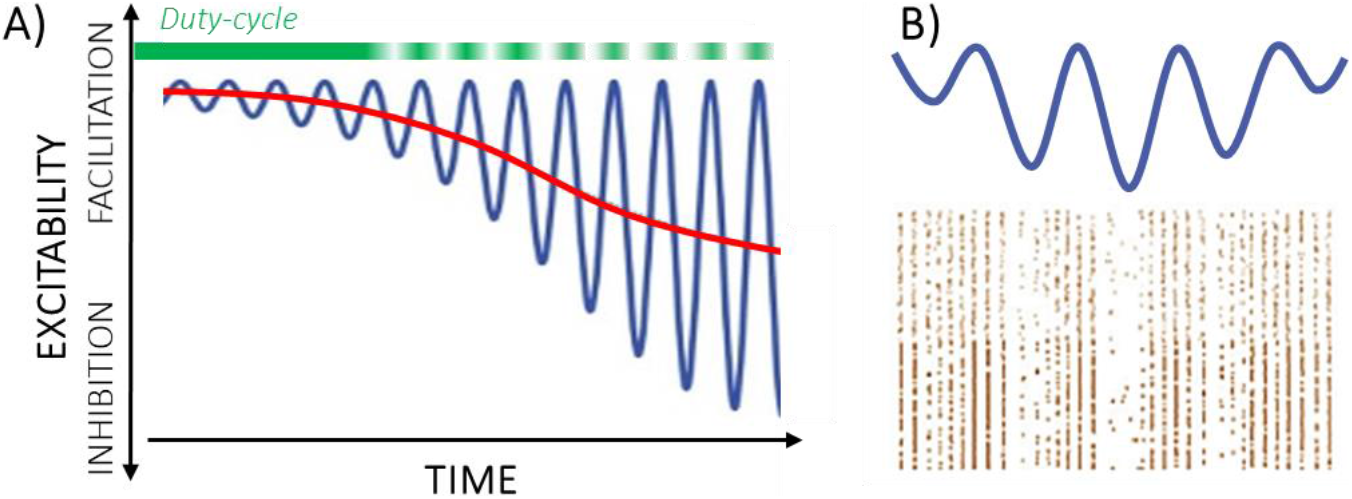
Negative and asymmetric alpha oscillations. **A)** An increase in alpha amplitude is accompanied by a modification in the mean voltage level (blue trace: time-varying instantaneous voltage; red trace: mean voltage level), which shortens the duty cycle (green trace); adapted from Schalk (2015). **B)** Alpha-band oscillations (blue trace) modulate neuronal spiking activity (bottom); neural spiking is phasically reduced, i.e., pulsed inhibition is stronger when the amplitude of alpha oscillations is higher; bifrom Hyafil et al. (2015).

Furthermore, alpha oscillations may be “asymmetric” or “biased” (Hyafil, Giraud, Fontolan, and Gutkin, 2015; Jensen and Mazaheri, 2010; Schalk, 2015), meaning that alpha oscillations are not zero-mean. Rather, the mean of an alpha cycle varies with the amplitude of alpha oscillation, as shown by the red trace in **Figure 2A**. According to this view, the instantaneous voltage level may represent a more direct measure of functional inhibition than power and phase individually, as suggested by the function-through-biased-oscillations (FBO) framework (Schalk, 2015; Schalk, Marple, Knight, and Coon, 2017). A higher amplitude of alpha activity (in absolute value) corresponds to a more negative instantaneous voltage level, which leads to higher inhibition (Hyafil et al., 2015; **Figure 2B**). Such interpretation is supported by invasive neurophysiological recordings, showing both a general decrease in neural firing rate during periods of high alpha power and a rhythmic relation between alpha oscillations and neuronal spiking (Haegens et al., 2011). FBO predictions are consistent with evidence showing the effects of alpha phase only when alpha power is high (Cohen and Van Gaal, 2013; Mathewson et al., 2009).

Concisely, negative asymmetric alpha oscillations represent a key element in inhibition.

## 3. ALPHA INHIBITION THROUGH ALPHA-GAMMA CROSS-FREQUENCY INTERACTIONS

Existing evidence (Bahramisharif et al., 2013; Jensen et al., 2014; Roux et al., 2013; Spaak et al., 2012) and current theories on the functional role of the alpha band (Bonnefond et al., 2017; Jensen and Mazaheri, 2010) suggest that the pulsed inhibition caused by alpha oscillations, which shortens the duty cycle, occurs by means of cross-frequency interactions with the gamma band. Importantly, although gamma activity is often associated with stimulus processing (Fries et al., 2007; but see Ray and Maunsell, 2015), alpha-gamma interactions have been consistently recorded during rest and prestimulus windows (Bahramisharif et al., 2013; Osipova et al., 2008; Spaak et al., 2012; but see Ray and Maunsell 2015), suggesting their crucial role in shaping the perceptual outcome (Ni et al., 2016a; van Es and Schoffelen, 2019).

As shown in **Figure 3**, the relationship between alpha and gamma oscillations is regulated both by amplitude-amplitude coupling (AAC), i.e., alpha power increase associated with gamma decrease, and by phase-amplitude coupling (PAC) interactions, i.e., gamma power nested within the alpha phase (Spaak et al., 2012). In line with the asymmetry in alpha oscillations, the stronger the alpha amplitude, the stronger the PAC (Osipova et al., 2008); high-amplitude alpha oscillations, accompanied by a general decrease in alpha voltage level, can lead to a stronger phasic modulation of gamma power compared to low-amplitude alpha oscillations (Hyafil et al., 2015). Therefore, PAC describes the phasic suppression of macro-scale M/EEG gamma activity within an alpha cycle.

**Figure 3.**
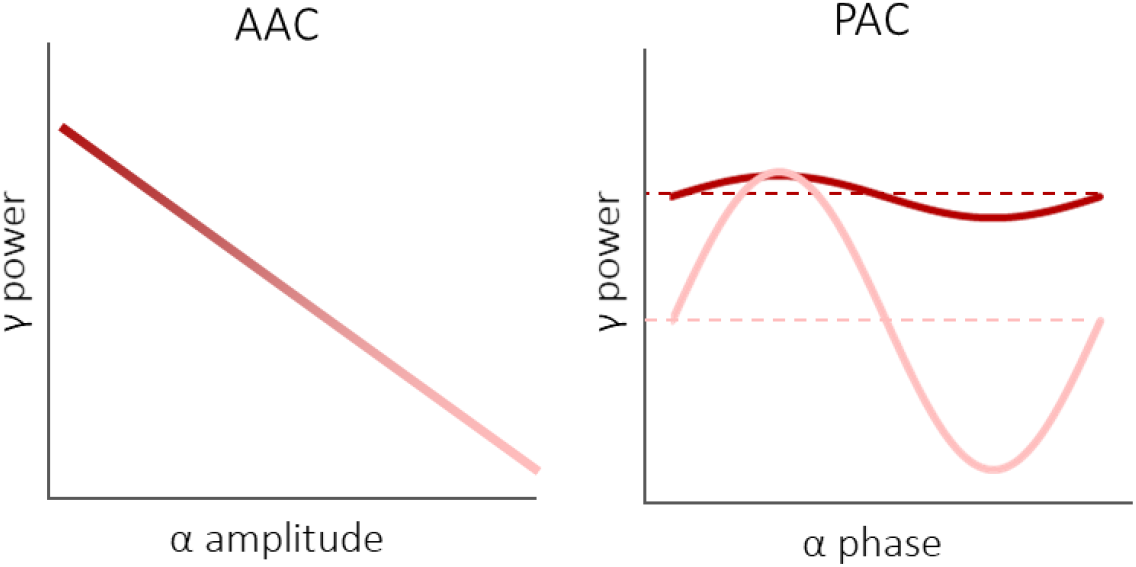
Schematic representation of alpha (α)-gamma (γ) cross-frequency interactions. *Left*: Amplitude-amplitude coupling (AAC); gamma power decreases as a function of alpha amplitude. *Right:* Phase-amplitude coupling (PAC) at different alpha amplitudes; when alpha is low (dark colour), the averaged gamma power (dashed line) is high with negligible influence of alpha phase, while when alpha is high (light colour), the averaged gamma power is low on average (dashed line) and is nested within the alpha cycle. Note that the maximum value of gamma power is the same for low and high alpha inhibition. Adapted from Hyafil et al., 2015.

In summary, we can conclude that alpha-gamma cross-frequency interactions are involved in ongoing oscillatory activity, in which alpha oscillations inhibit M/EEG gamma power.

## 4. ALPHA-MODULATED GAMMA OSCILLATIONS AFFECT RESPONSE PROBABILITY

The alpha-gamma AAC and PAC mechanisms are grounds to understand the inhibitory effects of ongoing alpha oscillations on perception: alpha can modulate the probability of response of gamma-oscillating neurons of sensory areas to incoming stimuli of different intensities (i.e., luminance or contrast level, considered to be proportional to the input to neurons and quantified in terms of depolarizing current; Tiesinga et al., 2004) and, consequently, the behavioural response probability.

Before taking into account alpha modulation of gamma activity, let us consider more deeply the probability of response of gamma oscillations. Gamma oscillations of small groups of neurons can be measured at the meso-scale level by means of invasive recordings, such as LFP, and more specifically, its local component (Kajikawa and Schroeder, 2011). **Figure 4A** shows the probability of response, according to the OPR model, when facing a wide range of input intensities in a condition without alpha inhibition: the neural response probability depends on a threshold (to be reached to induce a response), input intensity and amplitude of LFP gamma oscillations (Ni et al., 2016). If we consider a wide range of input intensities below a critical intensity (*a*), an incoming input will never induce a response (response probability = 0) in sensory neurons. For higher intensities, the neural response depends on the phase and amplitude of LFP gamma oscillations, giving rise to a response probability between 0 and 1. For even higher intensities, i.e., above a value (b), the stimulus will induce a response independently of gamma phase (response probability = 1). The resulting probability function of the neural response (from now on referred to as “single” probability of response) at different input intensities is represented in **Figure 4B** and mathematically defined as follows (formulas reported in **Table 1**). LFP gamma oscillations (GF) are defined in (1) as a sinusoidal function, while the single probability of response (PSR), independent of gamma frequency, is defined in (2), which is calculated as the ratio between the time window in which the input elicits a response (i.e., when the sum of gamma voltage and input intensity is higher than the threshold) and the wavelength of gamma. When considering a population of neurons (i.e., many small groups of neurons), the global probability of response is calculated by averaging the single response probabilities. Within a neural population, we can assume a certain degree of variability in the amplitude of LFP gamma oscillations (**Figure 4C**). Considering this variability to be normally distributed among neurons, the global response probability can be expressed as the weighted average of single response probability functions, as defined in (3). As shown in **Figure 4D**, the global response probability reveals a sigmoid trend. The link between the response of neural populations to the behavioural outcome is a highly complex subject, but, at least in simple visual perception tasks such as detection, we can assume the behavioural response probability to be proportional to one at the neural level (Britten, Shadlen, Newsome, & Movshon, 1992; Reynolds et al., 2000; Williford & Maunsell, 2019; but see Hara, Pestilli, & Gardner, 2014). Therefore, we consider that the global response probability function obtained from (3) may represent the psychometric function observed behaviourally (Britten et al., 1992), in which an observer’s performance in a visual detection task is related to the physical quantity of a stimulus, e.g., its intensity (Wichmann and Hill, 2001).

**Figure 4.**
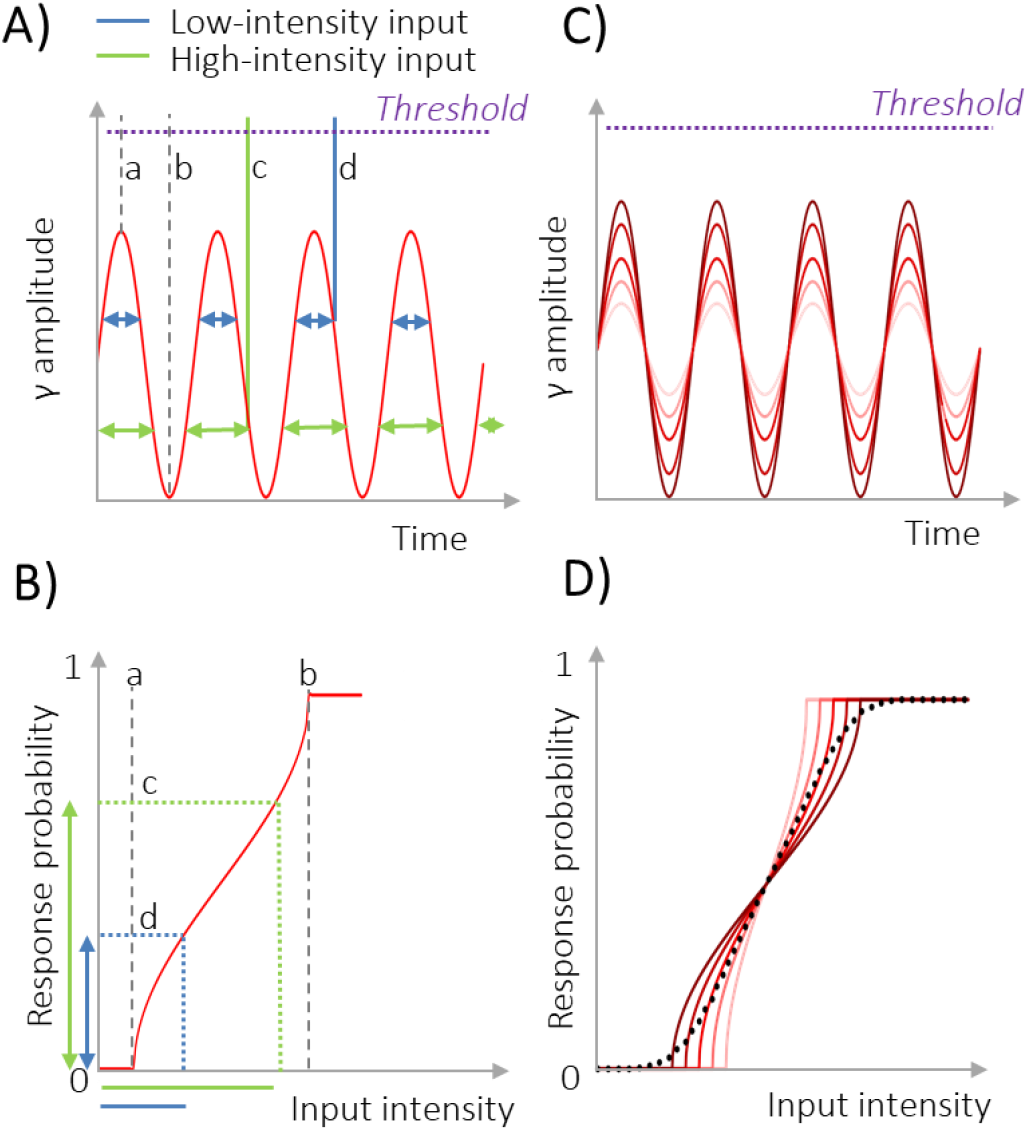
Neural response probability as a function of input intensity in a condition without alpha inhibition. **A)** Incoming inputs (*c:* high intensity, green; *d:* low intensity, blue) during LFP gamma oscillations of sensory neurons. Given a fixed threshold for response, double-headed arrows indicate the response probability (i.e., time windows during which incoming inputs may induce a response). **B)** Response probability as a function of input intensity at fixed threshold and fixed gamma amplitude, as defined in (3). Along the input intensity axis, *a* represents the minimum input intensity able to elicit neural response, *b* shows the minimum input intensity that determines a response independently of phase, and inputs between *a* and *b* indicate intensities inducing a phase-dependent response (probability between 0 and 1); *c* and *d* represent the high-and low-intensity inputs shown in (A). Along the response probability axis, *c* and *d* represent the response probability of the high- and the low-intensity inputs, respectively. **C)** Variability in the amplitude of LFP gamma oscillations within the population of sensory neurons leads to **(D)** variability in response probability functions. The mean of all response probability functions, weighted for their distribution in the neural population, results in a sigmoid function that represents the global response probability function (dotted line).

**Table 1.**

|  | Without alpha inhibition | With alpha inhibition |  |
| --- | --- | --- | --- |
|  |  | b.1 Gamma amplitude suppression | b.2 Gamma desynchronization |
| Alpha-gamma interaction | (1)<br>$G_F(t) = G \sin\left(\frac{2\pi}{L_G}t\right) + G_M$ | (4)<br>$A_F(t) = A \sin\left(\frac{2\pi}{L_A}t\right) + A_M$ | |
| | | (5)<br>$G_F(t) = K(A_F(t) - Min) \sin\left(\frac{2\pi}{L_G}t\right) + A_F(t)$ | (8)<br>$G_F(t) = G \sin\left(\frac{2\pi}{L_G}t\right) + A_F(t)$<br>(9)<br>$S_F(t) = \frac{A}{A + A_M - Min} \sin\left(\frac{2\pi}{L_A}t\right) + \frac{A_M - Min}{A + A_M - Min}$ |
| Single probability of response | (2)<br>$0 < G_M < (G_M + G) < T$<br>$\cdot \quad T - (G_M + G) < I < T - (G_M - G),$<br>$P_{SR} = 0,5 + \frac{1}{\pi} \arcsin \frac{(I - T + G_M)}{G}$<br>$\cdot \quad I \leq T - (G_M + G),$<br>$P_{SR} = 0$<br>$\cdot \quad I \geq T - (G_M - G),$<br>$P_{SR} = 1$ | (6)<br>$\forall t_i:$<br>$\cdot \quad T - G_{Max} < I < T - G_{Min},$<br>$P_{SRi} = 0,5 + \frac{1}{\pi} \arcsin \left\{ \frac{I - T + A_F(t_i)}{K[A_F(t_i) - Min]} \right\}$<br>$\cdot \quad I \leq T - G_{Max},$<br>$P_{SRi} = 0$<br>$\cdot \quad I \geq T - G_{Min},$<br>$P_{SRi} = 1$<br>where:<br>$G_{Min} = (1 - k)A_F(t_i) + KMin$<br>$G_{Max} = (1 + k)A_F(t_i) - KMin$ | (10)<br>$\forall t_i:$<br>$\cdot \quad T - G_{Max} < I < T - G_{Min},$<br>$P_{SRi} = \left\{ 0,5 + \frac{1}{\pi} \arcsin \left[ \frac{I - T + A_F(t_i)}{G} \right] \right\} * S_F(t_i)$<br>$\cdot \quad I \leq T - G_{Max},$<br>$P_{SRi} = 0$<br>$\cdot \quad I \geq T - G_{Min},$<br>$P_{SRi} = \left\{ 0,5 + \frac{1}{\pi} \arcsin \left[ \frac{G_{Min} + A_F(t_i)}{G} \right] \right\} * S_F(t_i)$<br>where:<br>$G_{Min} = A_F(t_i) - G$<br>$G_{Max} = A_F(t_i) + G$ |
| | | (7)<br>$P_{SR} = \frac{1}{L_A} \int_0^{L_A} P_{SRi} dt$ | |
| Global probability of response | (3)<br>$P_{GR} = \frac{\sum_{j=1}^n P_{SRj} w_j}{\sum_{j=1}^n w_j}$ | | |

Crucially, the OPR allows an estimation of the probability of response when gamma activity is modulated by alpha inhibition. The alpha-gamma interaction involves several features. First, as is typically shown in raw LFP signals (Jia and Kohn, 2011), gamma activity is regulated by asymmetric alpha oscillations in a phase-dependent manner; therefore, gamma fluctuates around the alpha voltage level. Moreover, gamma is regulated by mechanisms that generate the AAC and PAC observed in M/EEG. The OPR models two different meso-scale mechanisms that regulate alpha-gamma interaction, which lead to distinct effects on the final behavioural response, as we will describe in detail in the following section.

In sum, these observations reveal that the OPR predicts the probability of response of gamma-oscillating neurons of sensory areas and therefore the probability of a behavioural response, which is mediated by alpha activity.

## 5. PROBABILITY OF RESPONSE IS SELECTIVELY MODULATED BY DISTINCT MECHANISMS OF ALPHA INHIBITION

Despite compelling evidence about the inhibitory role of alpha activity on visual perception and on cross-frequency interactions with the gamma band, little is known about the micro- and meso-scale mechanisms associated with ongoing alpha inhibition (Cohen, 2017; Hyafil et al., 2015; Sadaghiani and Kleinschmidt, 2016; Spaak et al., 2012). Indeed, the general relation between M/EEG features and lower-scale mechanisms is likely to be few to some rather than one to one: the same M/EEG oscillatory feature (e.g., alpha activity) may be generated by distinct processes (Cohen, 2017). For example, M/EEG power appears to be generated at lower-scale levels, both by changes in amplitude, i.e., the magnitude of an oscillation (Whittingstall and Logothetis, 2009) and in synchronization, i.e., the temporal alignment of the phase of oscillations (Makeig et al., 2002). Specifically, evidence from simultaneous LFP and EEG recordings of the visual cortex in behaving monkeys has shown that LFP amplitude and synchronization independently contribute to M/EEG gamma power (Musall et al., 2014).

Based on this evidence, the OPR models these two mechanisms in the alpha-gamma cross-frequency interaction to explore their effects on the probability of response. Importantly, according to the OPR, alpha-driven changes in amplitude and in synchronization of LFP gamma oscillations lead to distinct effects on the global response probability function and therefore on the behavioural outcome. Thus, a psychophysical approach that describes behavioural responses for a wide range of stimuli, i.e., the psychometric function, provides a powerful tool that may be exploited to disentangle the effects of the two phenomena in alpha-driven modulation of sensory processing.

According to the OPR, alpha modulates M/EEG gamma power by decreasing (a) the amplitude of LFP gamma oscillations and/or (b) the degree of synchronization among small groups of neurons.

### (a) Alpha reduces LFP gamma amplitude

An increase in alpha oscillations may induce not only a rhythmic decrease in the mean LFP gamma voltage level but also a decrease in gamma amplitude, which leads to a reduction in M/EEG gamma power (**Figure 5**, blue box). Similarly, the amplitude of fast oscillations has been shown to be increased by depolarizing currents that bring membrane potentials closer to the firing threshold (Bracci et al., 2003). With the same mechanism, alpha activity may phasically move the mean voltage of gamma oscillations away from threshold and may induce a phasic reduction of these oscillations, the mechanism of which is mathematically defined as follows. Alpha (*A*_*F*_) is defined in (4), and alpha-modulated gamma (*G*_*F*_) is defined in (5): LFP gamma oscillation is modelled as a sinusoidal function with a mean equal to alpha voltage and amplitude proportional to alpha voltage. In this way, the gamma phase is independent of the alpha phase, so that at each time point in the alpha wave, the gamma phase can assume any value. In (6), the probability of response of sensory neurons (*P*_*SR*_) is defined for each time point (*t*_*i*_), by combining (2), which calculates the single response probability for gamma oscillations, with (4) and (5), which include the modulation of gamma by means of alpha. The single probability of response of neurons over the entire alpha cycle for each input intensity is calculated as the sum of response probabilities at each time point *t*_*i*_ with time interval *dt*; with *dt* close to zero, the calculation becomes more precise and is expressed by (7). As defined above, when considering a population of neurons, the global response probability (*P*_*GR*_) is expressed as the weighted average of single response probabilities (3) and results in a sigmoid trend.

**Figure 5.**
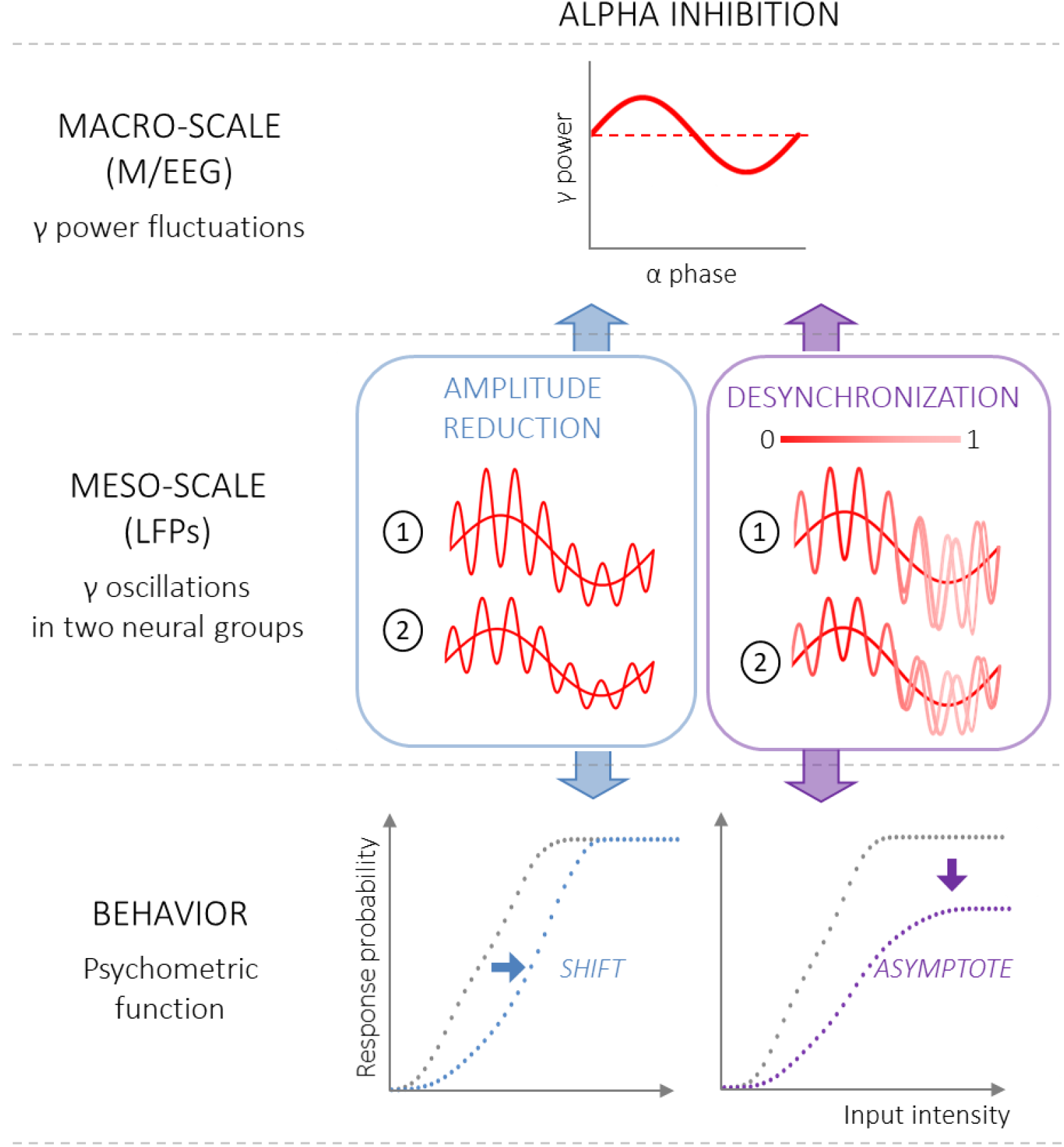
The OPR predictions associated with alpha inhibition described at different scales. ***Macro-scale*** *(top row):* gamma power, as measured by M/EEG recordings, is modulated by alpha inhibition by means of AAC and PAC mechanisms shown in Figure 3: M/EEG gamma power is lowered on average and is nested within the alpha phase. ***Meso-scale*** *(middle row):* two illustrative groups of sensory neurons, as measured by the LFP, oscillating in gamma at variable amplitudes and modulated by alpha activity. During alpha inhibition, the modulation of gamma power fluctuations may arise either from a phasic suppression of gamma amplitude (blue box) or from a phasic desynchronization of gamma oscillations (violet box; synchronization range as shown by colour bar). ***Behaviour*** *(bottom row):* different mechanisms of alpha inhibition lead to distinct effects on the psychometric function (dashed line: psychometric function without alpha inhibition). If alpha inhibition is associated with a phasic suppression of the amplitude of gamma oscillations, the OPR predicts a rightward shift of sensory threshold (blue line), while phasic gamma desynchronization is expected to give rise to a lowering of the upper asymptote (violet line).

A change in alpha activity modulates the psychometric function and determines a horizontal shift that can be calculated with a numerical simulation expressed by (6), (7) and (3). When alpha increases, the psychometric function is shifted rightwards, resulting in a worsening of visual performance, as shown by the blue trace in **Figure 5**.

In the literature on visual perception, a shift of the function is commonly described in terms of a contrast gain mechanism, arising from a divisive scaling of the input (Chaumon and Busch, 2014; Ling and Carrasco, 2006; Pestilli et al., 2007; Reynolds et al., 2000; van Boxtel, 2017). Interestingly, a contrast gain-like mechanism has been more associated with single-neuron activity than on the activity of the system as a whole (Kim et al., 2007), in line with the OPR. Moreover, a contrast gain-like mechanism has been suggested as the main functional process underneath sustained covert attention and adaptation, both from behavioural studies and single-unit neural measurements (Cameron, Tai, and Carrasco, 2002; Carrasco, Ling, and Read, 2004; Ling and Carrasco, 2006; Pestilli, Viera, and Carrasco, 2007; Reynolds et al., 2000).

### (b) Alpha reduces LFP gamma synchronization

Alpha-driven gamma power modulations may otherwise arise from phasic desynchronization of LFP gamma oscillations (**Figure 5**, violet box). In this case, the alpha-induced rhythmic decrease in the gamma mean is associated with a decrease in synchrony. The degree of neural synchronization is expressed in a range between 0 and 1, where 0 means complete gamma desynchronization among neurons and consequent zeroing of the global response despite the actual response of sensory neurons. This definition is in line with several measures adopted to quantify neural synchrony in the literature on functional connectivity, which rely on the concept of cross-correlation (e.g., the coherence coefficient; Bastos & Schoffelen, 2016). Here, alpha (*A*_*F*_) is defined in (4), alpha-modulated gamma (*G*_*F*_) is defined in (8), and synchronization is modelled in (9), as follows. The synchronization factor (*S*_*F*_), as expressed in (9) and shown in **Figure 6**, is modelled to be proportional to the alpha oscillation defined in (4): when the alpha voltage is at the maximum, the synchronization factor is 1; when the alpha voltage is at the minimum, the synchronization factor is at its minimum, and its value at each time point of the alpha wave is proportional to the gap between the alpha voltage and absolute minimum voltage.

**Figure 6.**
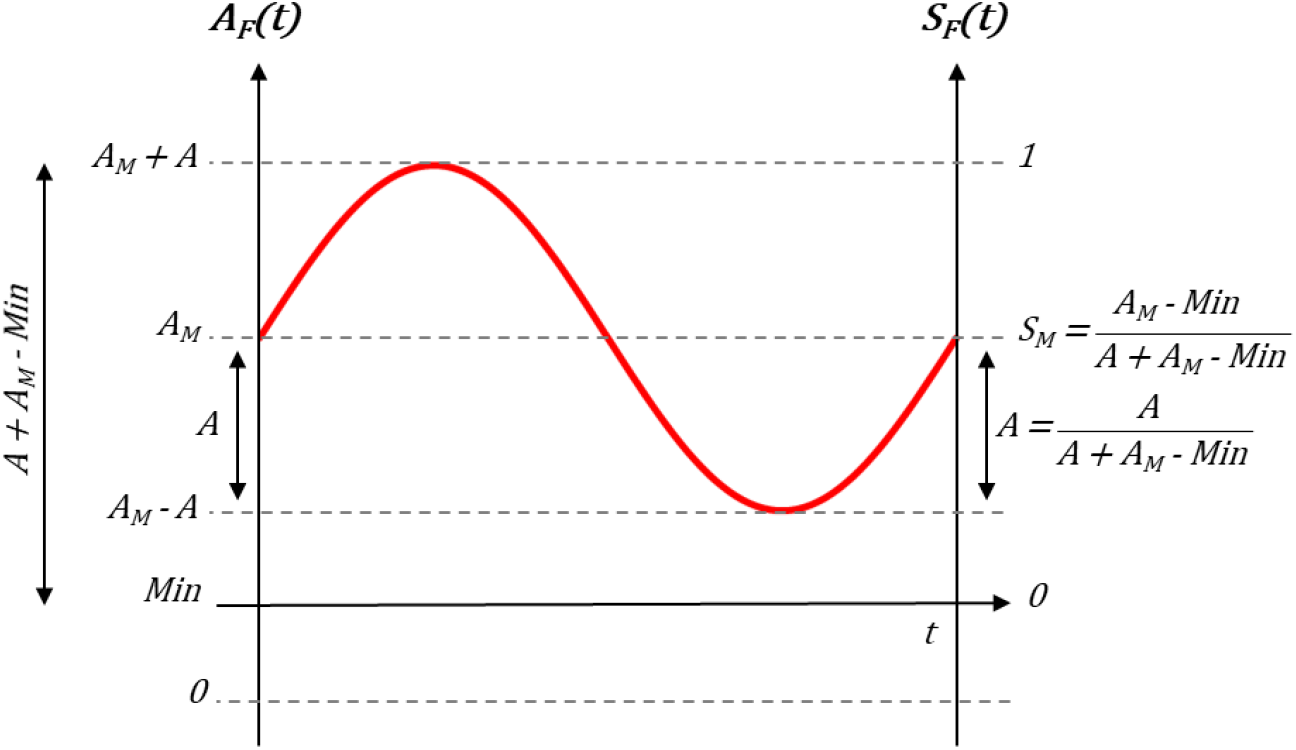
Synchronization factor. The synchronization factor (*S*_*F*_*(t)*) is defined as a sinusoidal function synchronized with *A*_*F*_. *S*_*F*_*(t)* is 1 when *A*_*F*_*(t)* is maximal (i.e., *A*_*F*_*(t)* = *A*_*M*_ + *A*), while 0 on the SF axis corresponds to the minimum voltage (*Min*) on the *A*_*F*_ axis. Therefore, interval 1 on the *S*_*F*_ axis corresponds to interval *A*_*M*_ + *A* – *Min* on the *A*_*F*_ axis, making the following proportions valid:

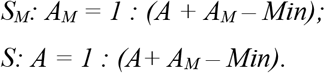

Postulating that the LFP gamma phase is independent from the alpha phase, the single probability of response is defined for each time point (*t*_*i*_) in (10), by combining (2) with (4) and (8), multiplied by the synchronization factor (9). Then, the single (*P*_*SR*_) and the global (*P*_*GR*_) probabilities of response are calculated in (7) and (3), respectively, consistent with that described in the previous paragraph. A numerical simulation of the psychometric function expressed by (7) and (10) results in a sigmoid trend. As shown by the violet trace in **Figure 5**, an increase in alpha activity associated with phasic gamma desynchronization leads to a worsening of visual performance, in this case by lowering the upper asymptote.

In the literature on visual perception, a modification in the upper bound of the psychometric function is referred to as the response gain mechanism, which mainly affects performance (as well as the neural response) to high-intensity stimuli because the modulation is proportional to the response (Chaumon and Busch, 2014; Reynolds et al., 2000). Intriguingly, neural synchronization has been proposed as a candidate mechanism for response gain (Buia and Tiesinga, 2006; Fries et al., 2001; Kim et al., 2007; Reynolds and Chelazzi, 2004). In previous studies investigating the relationship between attention and visual perception, the response gain mechanism has been related to transient and exogeneous attention (Ling and Carrasco, 2006; Pestilli et al., 2009).

In summary, the OPR predicts distinct effects of an alpha-induced LFP gamma amplitude decrease and gamma desynchronization on the psychometric function, i.e., a rightward shift of the curve and a lowering of the upper asymptote, respectively.

## DISCUSSION AND FUTURE DIRECTIONS

In the present work, we modelled the behavioural outcome of distinct meso-scale mechanisms that may be associated with the same M/EEG feature, i.e., ongoing alpha oscillations and cross-frequency alpha-gamma interactions. The OPR is intended as a simplification of the relationship between neural mechanisms and behaviour, and future studies combining simultaneous invasive and non-invasive recordings, as well as the development of computational models, are needed to test the OPR.

If applied in the context of non-invasive recordings, the OPR exploits the psychometric function estimation to make predictions on the meso-scale neural mechanisms involved in ongoing activity. Such an approach is not new in the literature on visual perception; indeed, the psychometric function estimation has proven to be extremely valuable for studying the effects of attention on visual perception, such as for testing for contrast gain *versus* response gain mechanisms (Cameron et al., 2002; Herrmann et al., 2010; Pestilli et al., 2009, 2007; Pham and Kiorpes, 2019; Reynolds et al., 2000; van Boxtel, 2017; Wunderle et al., 2015). Considering that attention has been related to ongoing alpha oscillations, we highlight that potential benefits could arise from extending this approach to the study of the effects of ongoing activity, revealing the existence of different processes associated with the same macro-scale phenomena. To our knowledge, only one study investigated the effects of prestimulus alpha activity on perception by relating the ongoing alpha power with changes in the psychometric curve as a function of stimulus intensity (Chaumon and Busch, 2014). Chaumon and Busch (2014) observed that higher prestimulus alpha power was associated with a decrease in the upper asymptote, which has been interpreted as reflecting a response gain mechanism. The OPR may extend findings from Chaumon and Busch (2014), allowing suggestions on the neural mechanisms associated with selective modifications of the psychometric function. Specifically, in this case, alpha power fluctuations could be interpreted as associated with gamma desynchronization in visual areas.

The OPR may shed light on the question of whether ongoing alpha rhythm represents a unitary phenomenon in M/EEG studies, for example, to explore if the same mechanisms are involved in spontaneous variations of ongoing alpha activity and when it is experimentally modulated (e.g., by means of spatial attention). While in both cases we can observe fluctuations in prestimulus alpha power across trials, the effect on the psychometric function of visual detection may differ, e.g., revealing a shift of the function or a lowering of the upper asymptote. If this is the case, then the OPR allows specific predictions on the possible mechanisms involved. However, the relationship among attention, ongoing alpha oscillations and changes in the psychometric function has not been directly addressed thus far. Establishing the relation between ongoing oscillations and visual perception in different experimental conditions is a critical matter for future research, and we argue that valuable insights may arise from combining the psychophysical approach of studies on visual attention and research on the effects of prestimulus oscillatory patterns.

In conclusion, while arising from the existing literature in the field of alpha oscillations on visual perception, the OPR may not only be extended to other perceptual domains but also encourage a fruitful discussion about the relationship between neural activity across different scales, i.e., meso- and macro-scales, and behaviour, bringing together efforts from several methodological and theoretical perspectives.

